# The DocMaps Framework for representing assertions on research products in an extensible, machine-readable, and discoverable format

**DOI:** 10.1101/2021.07.13.452204

**Authors:** Gary S. McDowell, Jessica K. Polka, Tony Ross-Hellauer, Gabriel Stein

## Abstract

Peer review of a research product varies widely depending on the publishers and platforms involved in the process. As scholarly publishing is disrupted by new innovations, peer review processes become more heterogeneous, placing an increasing burden on the researcher in understanding how they can communicate their scholarship. New ways to model such processes, and increase transparency, trust, and experimentation in scholarly publishing are needed. Many are emerging but can tend to focus on the needs of creators, and not those of readers, funders, and the whole scholarly publishing ecosystem. They may not place focus on representing editorial practices in ways that can be reliably aggregated, surfaced, and queried; are often limited to traditional peer review processes; and cannot capture the full range of editorial practices and events needed to accommodate alternative publication, review, and curation models. To support researchers in a world of experimentation in scholarly publishing, we propose a machine-readable, extensible, and discoverable framework for representing and surfacing review and editorial processes. Working with a Technical Committee composed of interested parties by employing a modified Delphi Method, we developed initial guiding principles and proposals towards an object-level editorial metadata framework compatible with a broad range of possible futures for scholarly publishing. We present the results of this process with a proposal and example use cases for *DocMaps*, a framework for representing object-level assertions.

## Introduction

Journals and publishers carry out a variety of processes on research manuscripts, from peer review to plagiarism checks, with the aim of ensuring quality and integrity in the scientific literature. These processes, checks, and transformations are highly heterogeneous across publishers and disciplines, and are becoming more so as scholarly publishing is disrupted by new innovations, the open science movement, and the removal of barriers to entry. But not only is the process by which these events take place highly varied, it is also opaque (Klebel et al. 2020). This means that the highly-trusted process by which knowledge is verified and established is largely based on faith that the processes take place, rather than in demonstration that an article has been thoroughly vetted. This in turn means that readers can struggle to establish how best to interpret and critique what they are reading; or for authors how best to communicate their scholarship to those readers.

This combination of opacity with heterogeneity derives from a traditional publishing model in which curation occurs before publication: authors select a journal that will curate their article first, prior to its publication and review. These three processes (curation, review, publication) could actually occur at different locations (e.g. a preprint server; a review service) and in a different sequence. But by prioritizing curation, with all processes taking place at that journal, the processes applied to a document are likely to be applied within a localized, and relatively closed, community of actors within a particular publishing platform. There are numerous experiments and platforms focusing on a “Publish-Review-Curate” model (Stern and O’Shea 2019) and it stands to reason that in a world where content is increasingly published first, then reviewed and curated, many different models for conducting this review will flourish, and experiments in review be undertaken. While this system may involve more transparent processes, it is still heterogeneous, and reviewers will need to be able to publish their reviews in a way that can be discovered and surfaced by publishers and aggregators. Publishers and aggregators will need ways to discover, normalize, and display reviews from multiple sources and taxonomies in consistent ways that readers can rapidly understand and contextualize.

There is a need to increase transparency about how scholarly publishing is undertaken. The COVID-19 pandemic, with its requirements for rapid review and an explosion of pre-printing, shows the urgency of this need.

While it may be tempting to suggest that there should be a standard way to undertake processes of publication, review and curation, we propose that what the current ecosystem needs instead is a framework that defines some general principles and best practices, while allowing the ecosystem to experiment, grow and evolve as needed. To address this need, we set out to construct the DocMaps Framework for capturing valuable context about the processes used to create research products. This framework was designed to capture as much (or little) contextual data about a document as desired by the publisher: from a minimum assertion that an event took place, to a detailed history of every edit to a document.

We started by identifying three key requirements for a framework for representing and surfacing object-level review and editorial events to create a healthy and trusted ecosystem:

- **Extensibility:** a wide range of editorial process events should be able to be represented, ranging from a simple assertion that a review occurred, to a complete history of editorial comments on a document, to a standalone review submitted by an independent reviewer.
- **Machine-readability:** assertions should be represented in a format that can be interpreted computationally and translated into visual representations.
- **Interoperability**: a single service should be able to interpret multiple taxonomies against the same criteria and arrive at the same interpretations.

The DocMaps project aims to assist the community by providing a framework to represent a broad range of editorial processes independent of the model used to define them; ultimately we aim to provide technical guides for implementing those requirements in a number of common formats.

The project took place in the midst of a rich ecosystem of initiatives related to models aiming to better describe and support peer review (see Appendix 1). Numerous recent initiatives have been positive developments in the space of peer review experimentation and transparency (Woolston 2019; Jones et al. 2020; “Peer Review Transparency,” n.d.; “Review Maps · KFG Notes” 2019). Such initiatives understandably tend to focus on the needs of the creators, and can be limited to traditional, even field-specific, peer review processes. They may not focus on representing editorial practices in ways that can be reliably aggregated, surfaced, and queried, or capture the full range of editorial practices and events needed to accommodate new publishing workflows in which reviews may be conducted by multiple parties. But they can be integrated into a framework like DocMaps to allow those who do, or wish, to use them to frame their contextual metadata around them. DocMaps aims to be a framework with a common way of describing editorial events, to which publishers of documents can add the components they can, or wish to, share.

Here, we describe our work in constructing the DocMaps Framework, using a modified Delphi Method (Linstone and Turoff 1975) to iteratively gather and prioritise the views of a Technical Committee (TC, see Appendix 2) comprising a broad-range of expertise and stakeholder viewpoints: parties interested in modelling object-level editorial processes; parties who may be willing to surface metadata as part of their infrastructure; and entities who create metadata as part of their editorial process. The outcome, also presented here, is a proposal for DocMaps, a common framework for representing object-level editorial processes.

In essence, the DocMaps Framework is composed of “DocMaps”, which are immutable assertions that describe an editorial event (or series of events). A “DocMap”:

- can describe multiple types of documents;.
- is asserted by the publisher of this document; and
- exists independently from another DocMap, but can reference other DocMaps.

In this way, a DocMap can describe a published paper, a preprint, a single referee report, an aggregate report, a recommendation, etc., allowing (for example) review services to curate evaluations from multiple sources in their reviews. At its simplest, a DocMap could simply describe the existence of an object:

~~~
**DocMap**
       {
       **createdOn**: timestamp // The time the DocMap was created
       **contentType**: string // The type of content (book, chapter, review)
       **content**: uri | optional // A link to the content the DocMap refers to
       **provider**: uri | optional // A link identifying the provider of the DocMap
(e.g. the journal)
       }
~~~

We here present our proposal for the consideration of the scholarly publishing community, providing two worked examples of use cases identified and described with the assistance of our TC.

## Method

The process of developing the DocMaps Framework chiefly relied on interacting with a Technical Committee (TC) composed of experts using a modified form of the Delphi Method (Linstone and Turoff 1975). Interactions with the TC were planned and executed by the Core Team (GSM, JKP, TRH, GS).

The TC was composed of parties interested in modelling object-level editorial processes; parties who may be willing to surface metadata as part of their infrastructure; and entities who create metadata as part of their editorial process. The list of permanent TC members involved throughout the process is described in Appendix 2.

The Delphi method is a systematic, interactive forecasting method which iteratively builds upon the combinatorial judgements of experts. The method seeks to increase the efficiency of face-to-face meetings by introducing additional structured group communications in between the meetings, in this case through the administration of strategically-designed surveys and requests for feedback. Time is then allowed for submission of responses, which are then anonymized and summarized by a facilitator. In this way, the method seeks to prevent group domination or “bandwagon” effects commonly encountered in face-to face meetings. In our version of the Delphi Method, members of the TC were not anonymous, and interacted with each other in regular (approximately monthly) one-hour meetings.

Following the traditional steps of the Delphi Method, the process was distinguished into four phases. The first exploratory phase (Phase I) allowed each TC member to contribute additional pertinent information, or make clear their initial reactions to the framing of the problem at hand. The next phase (Phase II) aimed to reach a reasonable consensus and shared understanding of how the group viewed the problem, including allowing discussions to clarify semantics or relative perspectives. The third phase (Phase III) sought to probe and explore areas of contention, and clarify outstanding issues, which in this case involved an initial draft proposal of the DocMaps Framework. The final phase (Phase IV) allowed us to present back to the group what had been heard, clarify any final questions and propose a final framework for publication.

In brief, the process involved the following steps:

1. An initial survey (Survey 1) prior to any face-to-face meetings, accompanying a description of the proposed project sent to participants who had previously agreed to take part in the TC (Phase I)
2. Meeting 1 (September 30th, 2020), where we presented the overall project, described the process, allowed group members to introduce themselves, and led a discussion of a summary of the results of Survey 1.
3. Survey 2 asked participants to suggest possible use cases for the DocMaps Framework, and asked for clarification on thoughts about an issue arising in Meeting 1, in order to reach consensus and clarify any arising issues (Phase II).
4. Survey 3 asked participants to vote on each proposed use case (summarized, standardized and anonymized) in order to help the group select two priority use cases to work up in the context of the DocMaps Framework.
5. Meeting 2 (October 29th, 2020) involved the TC picking two use cases, and breaking into two working groups to discuss the elements required to describe these use cases, and then gave data and their sources that would correspond to those elements.
6. Survey 4 proposed the first draft of the DocMaps Framework and asked for comments on the draft, as well as asking specific questions to probe and explore areas of contention (Phase III).
7. Meeting 3 (December 14th, 2020) extended this discussion of contentious issues into a face-to-face setting.
8. Meeting 4 (January 20th, 2021) brought real examples that could be applied to the use cases, to discuss the realities of implementing the proposed DocMaps Framework with the TC.
9. In Meeting 5 (February 10th, 2021), we presented a summary of the work undertaken and the final DocMaps Framework to be published in this work (Phase IV).

In addition to this work, we also discussed the DocMaps Framework proposal at an initial recorded webinar for our Co-Creation Community, a group of self-identifying interested parties who we are asking to give more feedback now and as the project progresses.

## Results

### Delphi Method with the Technical Committee

As described in the Methods, we first asked our TC to complete Survey 1 prior to any face-to-face meetings. This survey, accompanying a description of the proposed project, was designed to achieve the goals of Phase I of the Delphi Method by allowing each TC member to contribute additional pertinent information and make clear their initial reactions to the framing of the problem at hand.

Survey 1 asked:

1. Off the top of your head, what higher level categories of editorial or evaluative events should be supported by this framework?
2. What are the main aims, and hopes, of such a project for you and/or your organization, and the scholarly communications ecosystem more broadly?
3. What major milestones need to be reached between where we are now and the future you envision?
4. What are the major pitfalls that could disrupt the project?

In answer to question 1, TC members provided a list of editorial or evaluative “Events” that could be supported within the DocMaps Framework, which were summarized and are listed in Appendix 3. In summarizing the suggested events, we took care to highlight events listed in the peer review taxonomy developed by the International Association of Scientific, Technical and Medical Publishers (Jones et al. 2020).

The responses to question 2 were combined and summarized as shown in Figure 1.

**Figure 1:**
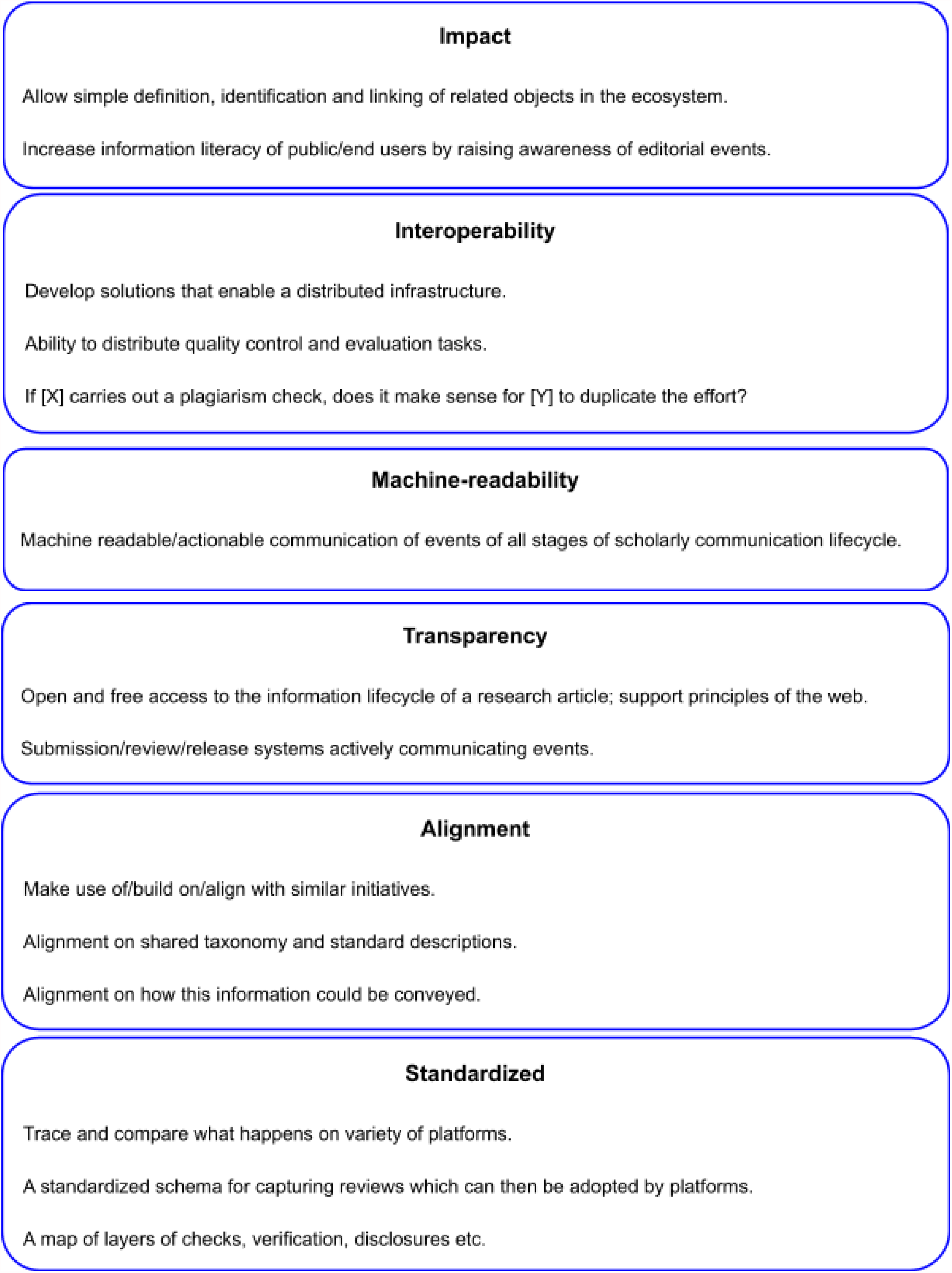
Summary of TC responses to the question: “What are the main aims, and hopes, of such a project for you and/or your organization, and the scholarly communications ecosystem more broadly?”

The responses to questions 3 and 4 were combined into Figure 2, as a timeline of the project was possible to combine with pitfalls that would arise at particular defined timepoints.

**Figure 2:**
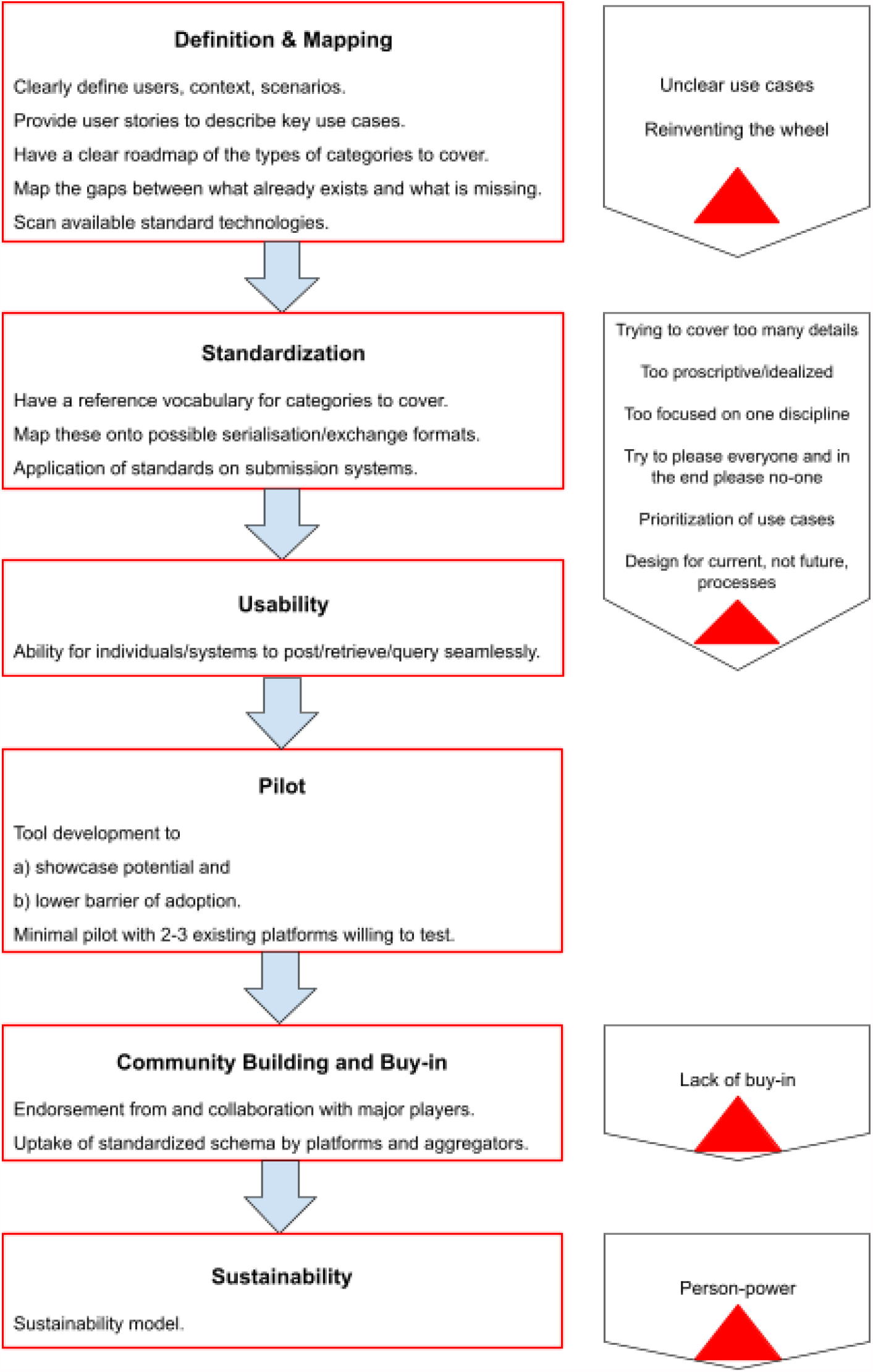
Combined summary of TC responses to the questions: “What major milestones need to be reached between where we are now and the future you envision?” (depicted on the left hand side) and “What are the major pitfalls that could disrupt the project?” (depicted on the right-hand side).

During Meeting 1, we led a discussion of a summary of the results of Survey 1. In the course of this meeting, a clear tension arose between the need to define events, and the desire to try to build a minimal pilot of something that could describe these events. In particular it was noted that other initiatives (such as the STM initiative (Jones et al. 2020)) were working on defining taxonomies, and that attempting to define every single possible event, and achieve consensus on these definitions, would be difficult and not a desired task for DocMaps.

Another issue that arose through the course of the meeting was clearly articulated by one participant as a collective issue in defining an event, and in particular the “status” of an event, and its perception in the community, or its “standing”. For example, an event can give an output (the “status” of the event) that may then be subjected to an assessment by a human, such as an editor (or, someone who makes an evaluation of its “standing”). This has previously been described extensively in discussions of preprints (Neylon et al. 2017).

We next asked the TC to provide responses to Survey 2 in order to reach consensus and clarify thoughts on the particular arising issue of status *vs*. standing (Phase II) and to provide suggestions for use cases that could be satisfied by the DocMaps Framework, in order to start working towards a proposal for a pilot.

Survey 2 asked:

1. Please give us at least 1 (maximum 3) use cases that you would like us to consider to ultimately work on to define Doc Maps.
  a. Feel free to describe the use case in as much detail as you would like; and cases can be as general or specific as you would like.
  b. You can make use of this list of possible events you provided to us.
  c. As a prompt, you may consider and expand upon use cases suggested previously.
  d. Please use the agile “user story” format of:
    i. “As a [type of user], I want [goal/function], so that [reason].”
2. In [Meeting 1], [PARTICIPANT] highlighted the question of the characterization of an event, particularly commenting on the status of an event (i.e.: what happened in the event with an objective measurement) *vs*. the standing of an event (i.e.: how the event is interpreted which is a subjective judgement (see (Neylon et al. 2017)). Our stance is that Doc Maps should list, as closely as possible, objective events, and leave subjective judgements and interpretations to others (This includes subjective judgements themselves being listed as events). Do you agree or disagree with this stance? Please give us your thoughts.

In response to question 1, a number of use cases were identified. The full list is provided in Appendix 4. The results were collated and then sent back to the TC, asking them to vote on which use cases they wished to prioritize (see Survey 3 below).

In general, respondents agreed with the stance in question 2. They noted that this stance would be likely to help build trust with organizations, and that as little room as possible should be left for interpretation and judgement. One example provided was that it would be correct to convey the event of e.g. an accept/reject decision, but not to convey how that decision was made.

A potential issue raised was that in service of constraining the framework to only objective events (or state changes), it would be hard to define such events, with suggestions that the group should try to specify some standard definitions. For example, in theory, excluding changes in standing could exclude recording publication or curation in journals, since they apply the value judgement as to whether a paper is worth the attention of their readers.

This concept was also articulated by TC participants as being potentially difficult for readers to understand, especially as separation of status from standing may not be trivial. For example, if a document was subjected to a plagiarism check, which was then signified through a badge (a change in status), that assumes and conveys to the reader that the article is plagiarism free (its standing), as per the standards and policies of the entity carrying out the check and providing this interpretation.

Survey 3 asked participants to vote on use cases proposed in Survey 2 (summarized, standardized and anonymized) in order to help the group pick 2 use cases to work up in the context of the DocMaps Framework.

Three use cases received the same, highest number of votes:

1. READER of a published article: wants to KNOW THE THOROUGHNESS of the peer review process of the article to BETTER DISTINGUISH trustable papers and MAKE A DECISION of further reading/paying to read/citing/sharing/etc.
2. JOURNAL: wants to PUBLISH its own reviews and the reviews transferred from another journal or peer review service alongside the published paper.
3. PREPRINT SERVER: wants to IDENTIFY AND SELECT third-party reviews to DISPLAY them alongside the associated preprint.

In Meeting 2, the TC agreed on two use cases, by combining the READER and JOURNAL use cases into a single use case, keeping the PREPRINT SERVER use case as stated. This followed from a discussion on the work that would be required in the READER use case of defining “thoroughness”, which was viewed as ambitious, and by some TC participants as desirable, but overall probably a significant effort which could distract from the goal of designing a simple pilot use case.

The TC then separated into two working groups during the meeting to discuss the elements required to describe these use cases, and identify data (and their sources) that would correspond to those elements. These results are described in Appendix 5.

In summary, the READER/JOURNAL group discussed elements that readers might want to see, or journals might want to communicate, to promote trust. This again brought up the issue of “status” *vs*. “standing”. Participants identified that there were many elements to draw from in the STM taxonomy (Jones et al. 2020). Participants discussed providing more transparency around quality control actions that journals may already do that are not communicated well at all, e.g. peer review of code. Participants also discussed the issue of expertise of reviewers (the field someone is in and how they contribute), and how difficult it would be to have consensus in describing this, perhaps needing the development of a taxonomy.

The PREPRINT SERVER group discussed the difference between accepting materials from a review service because they meet certain qualifications and agree to transmit certain pieces of information *vs*. allowing anyone to send in a review: what information you would need to determine whether to accept or reject displaying the review? The group discussed the pre-approval route and explored how the review would be transmitted, what format it would use, and how that would affect the different types of data that would be needed. There followed a discussion about all the types of data that would be needed, one being the structured full text for the review, for example allowing you to extract scores or ranking. As there is no accept/reject decision in the preprint case, participants suggested that taxonomies may need to be developed for scores and ratings.

The Core Team took the results from this discussion and expanded the use cases, categorizing and prioritizing elements for each use case. Using this, a first draft of the DocMaps Framework was created (see Appendix 6). To begin work on Phase III of the Delphi Method, to probe and explore areas of contention, and clarify outstanding issues, this draft was sent to the TC with Survey 4 with a request for comments on the draft:

1. Do you agree with the main concept for this structure?
2. Could this work in a better way?
3. What do you agree with?
4. What do you disagree with?

In response, TC participants sought clarification on two issues: the perceived tension between defining a taxonomy *vs*. a framework; and questions surrounding Versioning within the DocMaps Framework.

Participants perceived a tension between efforts to standardize or taxonomize editorial event information, and building a framework to represent the information in those standards and taxonomies (and beyond) in a way that can be interpreted and used in multiple formats, for multiple purposes.

Participants also questioned the logic of having ‘version’ as a value for ‘article type’, rather than as a separate element, pointing out that versions were iterations of an object rather than a new object itself. Would including version as a content type and a context introduce confusion? Would pre-publication revisions be classified as versions?

In order to explore these questions further, they were discussed in face-to-face Meeting 3. Participants discussed issues surrounding the location of metadata about content (e.g. a document) and about related objects (e.g. a review of that document), with each having its own DocMaps describing metadata about its own object. There was an agreement about the need for an entity to be able to knit all of these assertions together, and also where to store this metadata. Whether other organizations were already carrying out some of these functions, and how DocMaps differs, was discussed, and in particular TC participants identified one key value of DocMaps being its ability to use an open and distributed infrastructure. The DocMaps Framework would provide a structure that anybody could then use, and any organization could push or pull that information to or from any other organization. The pushing and pulling of information was identified as a problem which could be bracketed at this stage and addressed in a later phase of the project.

When discussing versioning, there was general consensus about a version being an attribute rather than a content type. Participants suggested learning from the web infrastructure itself and linking from one version to another indicating a predecessor or successor by means of reference rather than value from one version to another. This would have the advantage of a single update not being a huge burden on whoever owns the DocMap for an object. Another discussion centered around what version reviews of an object would sit on, i.e. the version that is reviewed, or the version that results from a review. This raised the question of the technical difficulty of linking reviews to an object that “doesn’t exist”, e.g. a version of a manuscript that is not publicly available and was only seen by internal reviewers, editors, and the authors. This was resolved as needing to place attributes on links as part of the DocMaps Framework, to capture this information.

In Meeting 4, the TC took a practical look into how building the Framework would look, using concrete examples, to see what issues could be exposed in the practical creation of the DocMaps Framework. This work was based on the previously identified uses cases, namely:

1. A publisher captures context about a review of an article published in their journal; and
2. An independent review service notifies a preprint server about a review of an article on their platform.

A community-driven technology effort, Sciety (sciety.org) is developing an application to support communities reviewing preprints, and their code is available on the articles reviewed already, and this provides an opportunity to work through a document which is deposited as a preprint, reviewed by a community with these reviews identified on Sciety, then published on in a journal. These events could approximate the two use cases identified by the TC. Using this example, the aim was to work with the TC to map out the events, and lay out a rough code for the DocMaps for the use cases. In doing this, we hoped to be able to have the TC identify any issues with the creation of DocMaps relating to a research product.

The example we used was deposited as a preprint on bioRxiv (Dubois et al. 2020) and published in eLife (Dubois et al. 2021) with the review process described in Sciety (https://sciety.org/articles/activity/10.1101/2020.02.20.958025).

We asked, in each case, what *events* had taken place that we would want to record in its DocMap? And as events were identified, were there any aspects of those events that should be described by the DocMaps Framework? We also began to build the DocMap during the meeting for each case.

In the process, we also learned that the same article was described on Early Evidence Base (https://eeb.embo.org/doi/10.1101/2020.02.20.958025), an experimental platform being developed by EMBO Press and SourceData, combining artificial intelligence with human curation and peer-review to highlight results posted in preprints. The example was pointed out as being particularly useful due to its illustration of a properly distributed system.

An analogy was made that the DocMap for the final article was somewhat like a rearview mirror of a vehicle i.e. of a process that had already happened, whereas the preprint case was like a view through the vehicle’s side window i.e. a view of a process in progress.

On the subject of how reviews are carried out in practice, there was a discussion of two situations:

1. a version rejected by one journal after a round of review, then the same version is independently reviewed completely independently by a different journal
2. a version that is on e.g. Review Commons involving the same journals, the decisions are not as a result of independent review processes.

These raise questions about what the DocMaps look like, how these are all linked together, and the importance of thinking about how things are grouped. Both the scenarios of solicited review events, and peer or community review events need to be encapsulated, and can stand alone as events. But when an editorial judgement is made, some entity then has to wrap these things and say because of these events, an editorial judgement has been made. Both things are possible, but there is a tension between the actors in the ecosystem that need to make this possible. The journal will publish a summary of solicited reviews and the version that was used for that, and the decision made on this basis; but a community review service may make an independent review and recommendation, and then some other entity again could then take all these things, and make a further judgement, citing those.

The final stage of the TC process (Phase IV) took place in Meeting 5, where a summary of the work undertaken was presented back to the group, clarifying what had been heard, any final questions and proposing a final DocMaps Framework to be published in this work.

### The DocMaps Framework

The text of the DocMaps Framework, providing an opportunity to comment, can be found at https://docmaps.knowledgefutures.org/pub/sgkf1pqa.

## Summary

DocMaps is a framework for capturing valuable context about the processes used to create documents, in a machine-readable way that can be expressed and interpreted in multiple formats depending on the use-case of the reader of the DocMap. The framework is designed to capture any amount of contextual data about a document — from a minimum assertion that a process took place, to a detailed history of every edit to a document.

We intend DocMaps to be general enough to capture the process behind any document, but our initial focus is capturing evaluative processes surrounding research documents — primarily peer review processes conducted by journals or by services reviewing preprints.

## Core Concepts

### DocMaps

In addition to being the name of the framework, a **DocMap** is the name of the object used to capture information about a document. At its most basic, a DocMap consists of just a few pieces of information:

~~~
{
createdOn: timestamp // The time the DocMap was created
contentType: string // The type of content (book, chapter, review)
content: uri | optional // A link to the content the DocMap refers to
provider: uri | optional // A link identifying the provider of the DocMap (e.g. the journal)
}
~~~

Using just these three pieces of information, a publisher could make a simple assertion that an article exists.

~~~
{
createdOn: 07-07-1999T00:00:00Z
contentType: “article”
content: https://doi.org/10.1109/5.771073
provider: https://ieee.org
}
~~~

Of course, this assertion leaves much to be desired. Most DocMaps will contain more information, e.g. the date an article was published; which version of an article the DocMap refers to; *etc*. DocMaps accomplishes this by building and maintaining schemas for different **Content Types**.

### Content Types

Content types extend the basic DocMaps structure to add information needed to describe a specific kind of content.

In the article example above, the publisher can provide more information about the article by looking at the schema for a DocMaps Article (see below) and providing as many of the fields as they feel necessary. For example:

~~~
{
createdOn: 07-07-1999T00:00:00Z
contentType: “article”
content: https://doi.org/10.1109/5.771073
provider: https://ieee.org
contributors: [{name: “N. Paskin”, type: “author”}]
title: “Toward unique identifiers”
}
~~~

Although the initial set of content types will be limited, working with relevant communities will enable the development of many different types that support their use cases. For example: reviews; versions; translations; and discussions.

### Contexts

When you create relationships *between* DocMaps to describe a series of events, these relationships are called **Contexts**, and are defined by a **context key** that describes the type of relationship and the direction of the relationship. Like content types, we imagine that many different types of Contexts will eventually be available, such as:

~~~
// context key: [list of DocMaps]
reviews: [Docmap]
isReviewOf: [Docmap] versions: [Docmap]
isVersionOf: [Docmap]
translations: [Docmap]
isTranslationOf: [Docmap]
discussions: [Docmap]
isDiscussionOf: [Docmap]
…
~~~

Using Contexts and Content Types, you can describe almost any kind of editorial process. For example, let’s imagine that the publisher wanted to let you know that two versions of the article exist. They could add Version Contexts to the Article to show that they have published two versions of the article:

~~~
{
createdOn: 07-07-1999T00:00:00Z
contentType: “article”
content: https://doi.org/10.1109/5.771073
provider: https://ieee.org
contributors: [{name: N. Paskin, type: author}]
title: ‘Toward unique identifiers’
versions: [
     {
     createdOn: 07-07-1999T00:00:Z
     contentType: “version”
     content: https://doi.org/10.1109/5.771073v1
     }
     {
     createdOn: 07-08-1999T00:00:Z
     contentType: “version”
     content: https://doi.org/10.1109/5.771073v2
     }
}
~~~

But the flexibility of DocMaps means they don’t have to publish all this information at once. They could also publish a DocMap describing just the second version of the paper by creating a Version DocMap with an *isVersionOf* context containing an Article DocMap.

~~~
{
createdOn: 07-08-1999T00:00:00Z
contentType: “version”
content: https://doi.org/10.1109/5.771073v2
provider: https://ieee.org
isVersionOf: [
     {
     createdOn: 07-07-1999T00:00:Z
     contentType: “article”
     content: https://doi.org/10.1109/5.771073
     }
]
~~~

Because contexts can flexibly describe events in many different ways, we expect that conventions will arise relatively organically between providers and consumers of DocMaps for describing different editorial processes at different levels of complexity.

As you’ll see in the use cases below, we’ve used the Versions context to provide information about both published article versions and review revision rounds. If a provider didn’t need or want to report revision rounds, or if there was only one round, they could omit the Versions context. In all of these cases, the consumer of the DocMap can decide how much of the context they want to use depending on how they plan to use the DocMap.

## Initial Use Cases

The Technical Committee specified two first use cases for DocMaps and further prioritized the data elements needed to capture relevant data for those cases. Below, we describe example DocMaps for these cases and explain how the elements requested by the Technical Committee can be imputed from the DocMaps.

### 1. A publisher captures context about a review of an article published in their journal

In this example, a journal is describing a double-masked peer review of an article with two rounds of revisions. They do this by nesting a Review context within an Article Context. They then further nest two Version Contexts within the Review Context to describe multiple rounds of feedback.

~~~
{
contentType: “article”
content: https://doi.org/article/123
createdOn: 2020-08-16T00:00:00Z
provider: https://myjournal.org
title: ‘An article about something!’
contributors: [
      {
      name: “Liz Jones”
      id: https://orcid.org/0002-0002 role: “author”
      }
      {
      name: “Eric Mays”
id: https://orcid.org/0005-0001 role: “data visualization”
      }
]
datePublished: 2020-01-01T:00:00Z
versions: [
      {
      contentType: “version”
      content: https://doi.org/article/123v1
      date_submitted: 2019-12-20T00:00:00Z
      date_online: 2020-08-15T00:00:00Z
      ethics_statements: “This was conducted ethically.”
      ocmpeting_interests: “There were no conflicts of interest.”
      }
]
reviews: [
      {
      contentType: “review”
      content: https://doi.org/review/abcd
      createdOn: 2020-06-01T00:00:00z
      provider: https://myjournal.org
      decision_date: 2020-07-20T00:00:00z
      decision: ‘accept with revisions’
      contributors: [
       {
       name: “John Doe”
       affiliation: “Wassamatta U”
       roles: [editor, author]
       }
       {
       id: 12345
       roles: [reviewer]
       }
       {
       id: 23456
       roles: [reviewer]
       }
       ]
       identity_transparency: ‘double-anonymized’
       reviewer_interacts_with: [editor]
       review_information_published: [editor-identities]
       versions: [
       {
       contentType: “version”
       createdOn: 2020-06-15T00:00:00Z
       contributors: [
       {
       id: 12345
       roles: [reviewer]
       }
       {
       id: 23456
       roles: [reviewer]
       }
       ]
       }
       {
       contentType: “version”
       createdOn: 2020-07-10T00:00:00Z
       date_online: 2020-08-15T00:00:00Z
       contributors: [
       {
       id: 12345
       roles: [reviewer]
       }
      ]
     }
    ]
   }
  ]
}
~~~

### 2. An independent review service notifies a preprint server about a review of an article on their platform

In this example, a review service is describing a fully transparent review of a preprint article with links to the review report and author response. They do this by including a content field for the review object and filling out the author response and STM Association Taxonomy metadata to describe the process of the review.

~~~
{
contentType: “review”
content: https://doi.org/review/123
createdOn: 2020-08-01T00:00:00z
provider: https://myreviewservice.org
decision_date: 2020-07-20T00:00:00z
decision: “accept”
contributors: [
       {
       name: “Tricia McMillan”
       affiliation: “Maximegalon University”
       roles: [editor, author]
       id: https://orcid.org/0000-0000
       author_suggested: false
       },
       {
       name: “Zaphod Beeblebrox”
       affiliation: “Betelgeuse State College”
       roles: [reviewer]
       id: https://orcid.org/0001-0001
       author_suggested: true
       }
       {
       name: “Arthur Dent”
       affiliation: “BBC”
       roles: [reviewer]
       id: https://orcid.org/0002-0002
       }
       {
       name: “Ford Prefect”
       affiliation: “Pan Galactic Gargle Blaster Society”
       roles: [invited_reviewer]
       id: https://orcid.org/0002-0002
       }
]
author_responses: [{
       contentType: “version”
       content: https://doi.org/response/123
       date_online: 2020-07-31T00:00:00Z
       date_submitted: 2020-07-15T00:00:00Z
}]
identity_transparency: [all-identities-visible, opt-in]
reviewer_interacts_with: [editors, reviewers, authors]
review_information_published: [reviewer-identities, editor-identities,
review-reports-author-opt-in]
versions: [
       {
       contentType: “version”
       date_submitted: 2020-06-15T00:00:00Z
       date_online: 2020-07-31T00:00:00Z
       }
]
isReviewOf: [
       {
       contentType: “article”
       content: https://doi.org/preprint/123
       }
]
~~~

### Draft Content Type Schemas

These are initial drafts of content type schemas, and will be somewhat incomplete by design. They are intended to capture the information necessary to make the two initial use cases listed above possible.

#### Article

~~~
{
// Common DocMap fields
contentType: “article”
content: uri // A canonical link or DOI to the article, if it exists
createdOn: timestamp
provider: uri
// Article specific metadata
title: string
abstract: string
contributors: [Contributors]
datePublished: date
// Contexts
versions: [Version]
}
~~~

#### Review

~~~
{
// Common DocMap fields
contentType: “review”
content: uri // A canonical link or DOI to the report publication, if it exists
createdOn: timestamp
provider: uri
// Review specific metadata fields
decision_date: timestamp
decision: string
contributors: [Contributor] // Includes reviewers, editors, asked reviewers
author_response: uri
identity_transparency: string // STM Assoc taxonomy
reviewer_interacts_with: [string] // STM Assoc taxonomy
review_information_published: [string] // STM Assoc taxonomy
post_publication_commenting: [string] // STM Assoc taxonomy
// Contexts
isReviewOf: [Article] // A list of DocMaps of reviewed objects
}
~~~

#### Version

~~~
{
// Common DocMap fields
contentType: “version”
content: uri // A canonical link or DOI to the version of the referenced
object, if it exists
createdOn: timestamp
provider: uri
// Version specific metadata
date_submitted: timestamp
date_online: timestamp
ethics_statements: string
competing_interests: string
data_availability: string
contributors: [Contributor]
}
~~~

#### Implementation

Once we discuss and come to agreement on the initial schemas, we’ll turn to implementation. We’re using pseudocode in the examples above because we want DocMaps to support multiple formats, since publishers’ technological needs and capabilities are so varied.

### Case Study

Here, we present an example of how a real situation could be described using the DocMaps Framework. In the example presented below, a document was deposited as a preprint on bioRxiv and after a review process described in Sciety and on Early Evidence Base, was published in eLife.

#### 1. A publisher captures context about a review of an article published in their journal

~~~
**DocMap** {
       contentType: ‘article’
       versions: [
           0: {id: ‘10.1101/2020.02.20.958025’}
           1: {id: ‘10.7554/eLife.59907’}
       ]
       reviews: [{
           version: 0
           contributors: [
                {name: ‘Christopher Warren’, role: ‘reviewer’}
                {name: ‘‘, role: ‘reviewer’}
                {name: ‘‘, role: ‘reviewer’}
                {name: ‘Thorsten Kahnt’, role: ‘reviewing_editor’}
                {name: ‘Christian Büchel’, role: ‘senior_editor’}
       ]
          editorial_decision: ‘accept’
          …
       }]
    author_response {
         version: 1
         uri: ‘https://…‘
              …
    }
}
~~~

#### 2. An independent review service notifies a preprint server about a review of an article on their platform

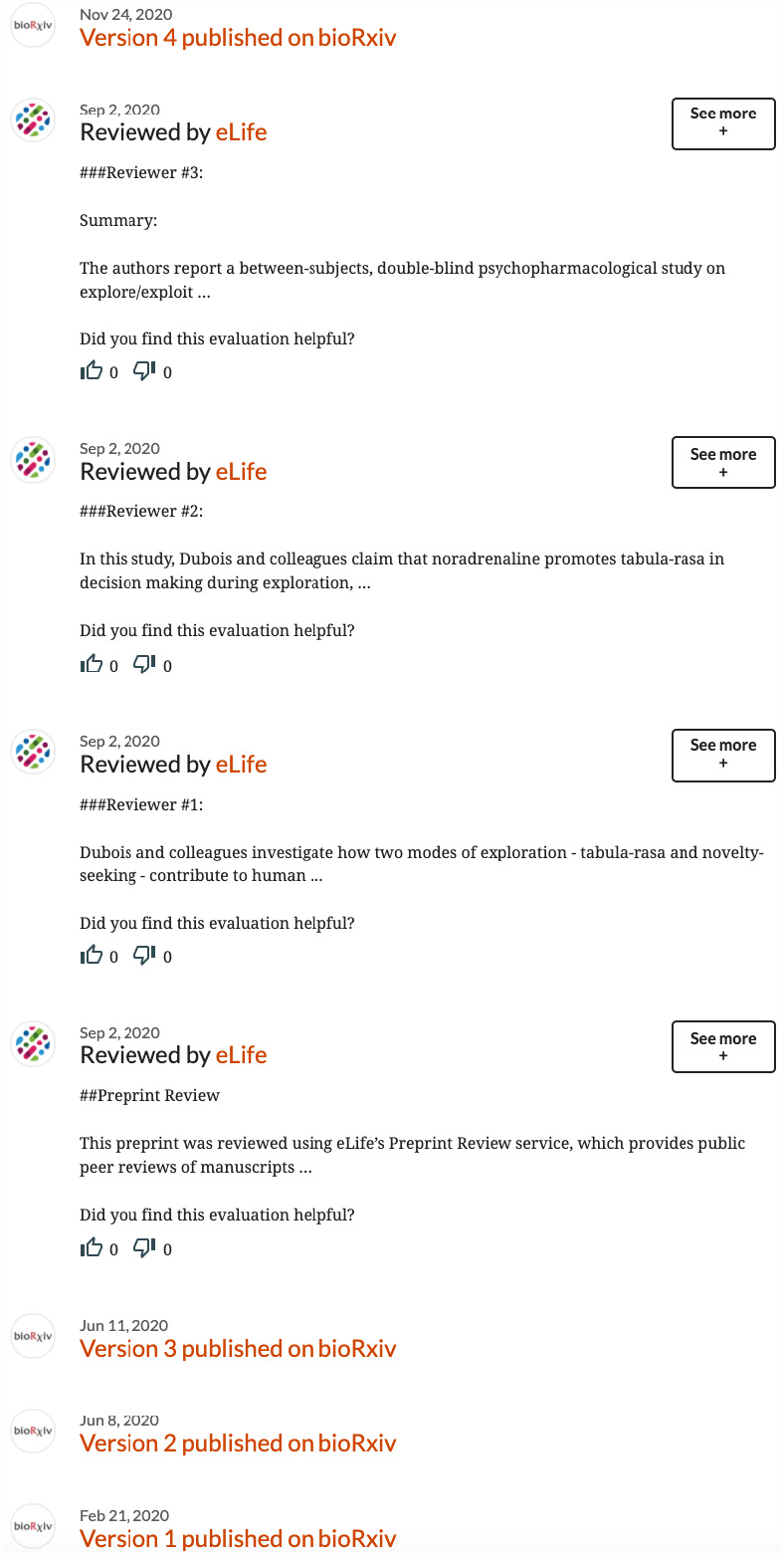

~~~
**DocMap** {
         contentType: ‘referee_report’
         id: ‘hyp.is/…’
         contributors: [{name: ‘‘, role: ‘reviewer’}]
         isReviewOf: ‘10.1101/2020.02.20.958025v3’
         …
}
**DocMap** {
         contentType: ‘referee_report’
         id: ‘hyp.is/…’
         contributors: [{name: ‘‘, role: ‘reviewer’}]
         isReviewOf: ‘10.1101/2020.02.20.958025v3’
         …
}
**DocMap** {
         contentType: ‘referee_report’
         id: ‘hyp.is/…’
         contributors: [{name: ‘‘, role: ‘reviewer’}]
         isReviewOf: ‘10.1101/2020.02.20.958025v3’
         …
}
**DocMap** {
         contentType: ‘review_aggregate’
         id: ‘hyp.is/…’
         contributors: [{name: ‘‘, role: ‘reviewer’}]
         isReviewOf: ‘10.1101/2020.02.20.958025v3’
         …
}
~~~

This describes one version of what the DocMaps could look like, with a DocMap for each review and the summary. These reviews are published individually to Hypothesis, so there’s another way that Sciety could theoretically describe these, if each version on Hypothesis had published an individual DocMap. They could curate those individual docmaps and produce a combined review DocMap, as shown below. Interestingly, this looks somewhat similar to the DocMap shown above for Use Case 1. However, it is possible that grouping reviews like this sends a signal about how the process was conducted, suggesting all reviews were part of a round of review stewarded by an editor.

~~~
**DocMap** {
         contentType: ‘review’
         id: ‘https://sciety.org/articles/10.1101/2020.02.20.958025‘
         isReviewOf: ‘10.1101/2020.02.20.958025v3’
         reviews: [
                 DocMap: {
                    id: ‘https://hyp.is/docmap/0.1101/2020.02.20.958025r1‘
                 }
                 DocMap: {
                    id: ‘https://hyp.is/docmap/0.1101/2020.02.20.958025r2‘
                 }
                 DocMap: {
                    id: ‘https://hyp.is/docmap/0.1101/2020.02.20.958025r3‘
}
                 DocMap: {
                    id: ‘https://hyp.is/docmap/0.1101/2020.02.20.958025s1‘
                 }
         ]
}
~~~

## Discussion

### Technical Limitations

As with any group selected for development of an idea, the initial proposal will be limited by the perspectives, experiences, assumptions and motivations of the people selected to be on the Technical Committee. It is for precisely this reason that we are proposing this Framework as an idea to the wider community, and soliciting feedback as part of our larger Co-Creation Community (C3) effort with a larger group of stake-holders. This process is open-invitation and aims to work with interested stakeholders towards implementation of DocMaps to suit their needs.

### Status vs Standing

A key tension that arose as part of this process was clarifying how, and indeed whether, DocMaps could list objective events without attaching subjective judgements and interpretations. In collectively discussing what “an event” in DocMaps is, we were able to articulate it in terms of the characterization of the objective elements of the event (its “status”) compared to its interpretation or subjective judgement (its “standing”). The view was that DocMaps would aim to transmit information about an objective measurement. The issue of representing the “status” of an event versus its “standing”, has been discussed previously with respect to preprints, with the authors pointing out that the issue becomes politicized as people mix these two concepts (Neylon et al. 2017). For example, the “status” of a preprint (e.g. its existence as a document on a server) is one thing, but the “standing” is then for the wider community to make a determination of how what that object means, for example in terms of the contribution of its contents to a body of knowledge. One outcome of this discussion was the agreement that it is not a goal of DocMaps to become a registry or accreditation agency for post-publication review services, for example. The aim is, as far as possible, to be content agnostic.

### Increasing Trust in Scholarly Publishing

One hope is that the information recorded in the DocMaps framework could be used to increase trust in scholarly publishing. In particular, we see a role in addressing concerns about misinformation with respect to preprints, and to clarify the status of retracted works, which have come to the fore during the COVID-19 pandemic. For example, an article funded by an organization with a particular agenda could currently be posted on a preprint server that does not undertake any quality control or screening, and used to spread disinformation. To most observers, including journalists and the general public, looking at the article on this server alone would not tell you that no quality control or screening has been undertaken, and it could have all the appearance of a legitimate article. If scholarly repositories surfaced information about editorial processes, the difference between an article (and its vetting process) on different servers could be readily apparent. Likewise, clear machine-readable ways of flagging retractions would make discredited works more readily identifiable as such, and even allow publishers to demonstrate that retractions of published papers can lead to improvements in the review process. DocMaps would make more data about editorial processes accessible, allowing researchers to systematically study processes and because the data would be public and interoperable, this could be done repeatedly and independently.

### Avoiding defining and prescriptive taxonomic definitions

There can be a temptation to be prescriptive as providers of information, and to attempt to define taxonomies for how information can be represented. DocMaps aims to be agnostic to these definitions, and to focus instead on recording the existence of the underlying editorial events that shape taxonomies and workflows. One publisher may not want to consider including reviews of objects that don’t exist publicly, and can ignore them. But another publisher could, and should be able to do so. We expect that publishers and consumers of DocMaps will describe and interpret the events provided according to their needs. The aim is to give people the data they need to make those judgements. This is a major difference between DocMaps and other schemas, taxonomies and formats: to allow the publication of information events in as flat a format as possible. The interpretation is done by the consumer of the events. In this way, we aim to ensure flexibility, and machine-readability, of contextual information for all types of uses, including non-traditional and even non-academic ones. We expect that conventions for publishing and interpreting DocMaps that can be used with specific taxonomies and workflows will arise, and intend to provide venues to support the development of these conventions.

## Conclusions and Future Directions

Using a modified Delphi Method (Linstone and Turoff 1975) we were able to iteratively gather and prioritise the views of a Technical Committee to construct a proposed common framework, DocMaps, to represent object-level editorial processes in a machine-readable, interoperable and extensible manner.

The DocMaps Framework describes editorial events through immutable assertions known as “DocMaps”, which:

- can describe multiple types of documents;.
- are asserted by the publisher of this document; and
- exist independently from another DocMap, but can reference other DocMaps.

In this way, a DocMap can describe a published paper, a preprint, a single referee report, an aggregate report, a recommendation, etc., allowing (for example) review services to curate evaluations from multiple sources in their reviews. At its simplest, a DocMap could simply describe the existence of an object.

Our proposal is for the consideration of the scholarly publishing community. Our future work involves gathering feedback on this proposal, with the assistance of a Co-Creation Community open to all interested parties, and [working with](https://docmaps.knowledgefutures.org/pilot) interested stakeholders to develop pilot implementations and experiments using the proposed DocMaps Framework. As a first example, we are working with EMBO’s [Early Evidence Base](https://eeb.embo.org/) and eLife’s [Sciety](https://sciety.org/) review curation platform, in conjunction with Cold Spring Harbor Lab’s [biorxiv](https://www.biorxiv.org/) and [medrxiv](https://medrxiv.org/) preprint servers, to pilot using DocMaps to represent and display distributed review events across platforms. These pilots will allow us to evaluate the utility and limitations of DocMaps in practice.

## Acknowledgements

We are grateful to the Howard Hughes Medical Institute for their support for this phase of the project. We would like to thank members of the Technical Committee for their participation in the process, and thank both members of the Technical Committee and the Co-Creation Community for their comments on this document prior to its publication.

## Appendix 1

### Other existing initiatives

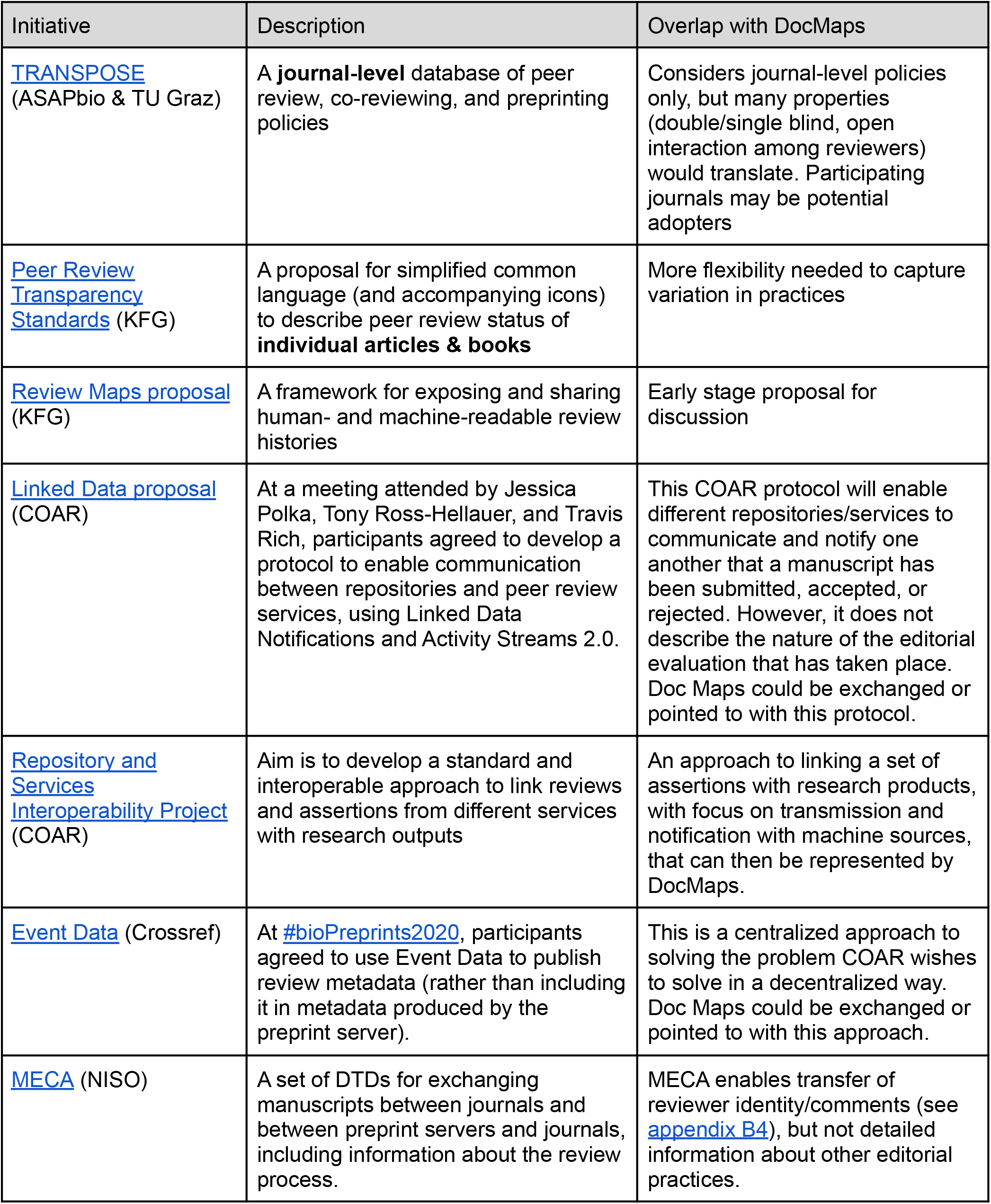
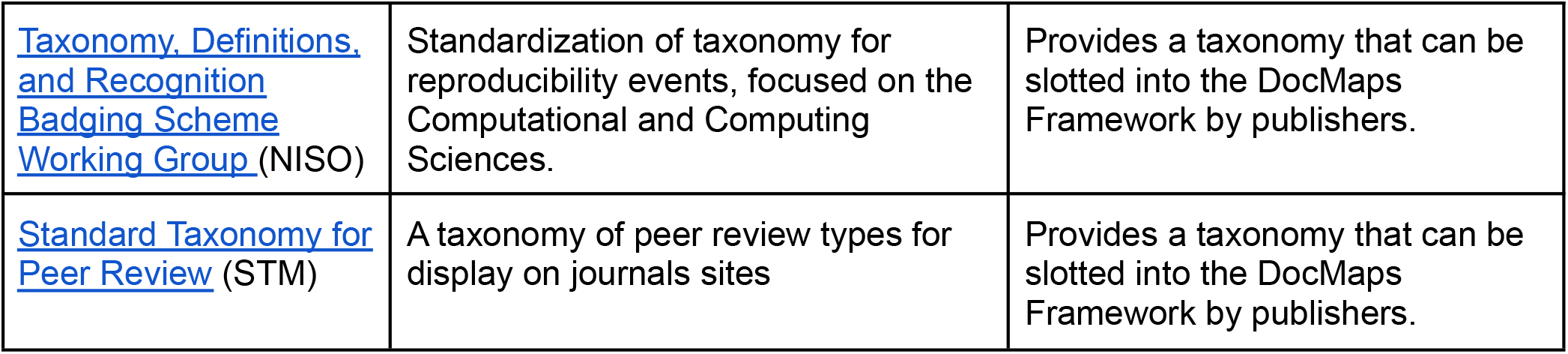

Appendix 1: Other existing initiatives
| Initiative | Description | Overlap with DocMaps |
| --- | --- | --- |
| TRANSPOSE<br>(ASAPbio & TU Graz) | A **journal-level** database of peer review, co-reviewing, and preprinting policies | Considers journal-level policies only, but many properties (double/single blind, open interaction among reviewers) would translate. Participating journals may be potential adopters |
| Peer Review Transparency Standards (KFG) | A proposal for simplified common language (and accompanying icons) to describe peer review status of **individual articles & books** | More flexibility needed to capture variation in practices |
| Review Maps proposal (KFG) | A framework for exposing and sharing human- and machine-readable review histories | Early stage proposal for discussion |
| Linked Data proposal (COAR) | At a meeting attended by Jessica Polka, Tony Ross-Hellauer, and Travis Rich, participants agreed to develop a protocol to enable communication between repositories and peer review services, using Linked Data Notifications and Activity Streams 2.0. | This COAR protocol will enable different repositories/services to communicate and notify one another that a manuscript has been submitted, accepted, or rejected. However, it does not describe the nature of the editorial evaluation that has taken place. Doc Maps could be exchanged or pointed to with this protocol. |
| Repository and Services Interoperability Project (COAR) | Aim is to develop a standard and interoperable approach to link reviews and assertions from different services with research outputs | An approach to linking a set of assertions with research products, with focus on transmission and notification with machine sources, that can then be represented by DocMaps. |
| Event Data (Crossref) | At #bioPreprints2020, participants agreed to use Event Data to publish review metadata (rather than including it in metadata produced by the preprint server). | This is a centralized approach to solving the problem COAR wishes to solve in a decentralized way. Doc Maps could be exchanged or pointed to with this approach. |
| MECA (NISO) | A set of DTDs for exchanging manuscripts between journals and between preprint servers and journals, including information about the review process. | MECA enables transfer of reviewer identity/comments (see appendix B4), but not detailed information about other editorial practices. |
| Taxonomy, Definitions, and Recognition Badging Scheme Working Group (NISO) | Standardization of taxonomy for reproducibility events, focused on the Computational and Computing Sciences. | Provides a taxonomy that can be slotted into the DocMaps Framework by publishers. |
| Standard Taxonomy for Peer Review (STM) | A taxonomy of peer review types for display on journals sites | Provides a taxonomy that can be slotted into the DocMaps Framework by publishers. |

## Appendix 2

### Technical Committee Membership

**Thomas Lemberger**, Embo Press

**Tony Alves**, NISO Manuscript Exchange Common Approach Standing Committee

**Laurent Romary**, INRIA

**Paul Shannon**, eLife

**Kathryn Funk**, Program Manager, PubMed Central, US National Library of Medicine, National Institutes of Health

**Richard Sever**, Co-Founder bioRxiv/medRxiv and Assistant Director CSHL Press, Cold Spring Harbor Laboratory

**Johanna McEntyre**, Europe PubMed Central

**Sowmya Swaminathan**, Head of Editorial Policy, Nature Research & Springer Nature

**Veronique Kiermer**, Public Library of Science

**Jennifer Lin**, Product Director, Meta at Chan Zuckerberg Initiative

**Martin Klein**, Los Alamos National Laboratory, Research Library

**Patricia Feeney**, Head of Metadata, Crossref

**Bhavani Shekhawat**, Software Engineering Team Lead, Elsevier

**Joris van Rossum**, Project Director, International Association of Scientific, Technical, and Medical Publishers

**Kathleen Shearer**, Executive Director, Confederation of Open Access Repositories (COAR)

**Caroline Webber**, Aries Systems Corporation Alumni:

**Bahar Mehmani**, Reviewer Experience Lead, Elsevier

**Damian Pattinson**, eLife

## Appendix 3

### List of Events suggested in Survey 1

STM taxonomy events (Jones et al. 2020) are marked in blue.

#### Pre-submission

- Registration
- Various parts/components/artifacts of the scholarly object

#### Type of Document

- Peer-reviewed-article
- Referee-report
- Editor-report
- Author-comment
- Community-comment
- Aggregated-review-documents
- Peer-review-report
- Major changes to article type, title, author lists, funders
- Version retraction
- Version withdrawal

#### Identity transparency (both editorial and review)

- Triple blind
- Double blind
- Single blind
- Visible

#### Submission/Deposition

- Date of submission
- pre-print service
- Journal
- conference submission system
- institutional repository
- Request for review (by anyone against any review entity) w/ corresponding response
- Number of reviewers
- Types of reviews

#### Review

- Date
- Reviewer
- Recommendation
- review text
- Author responses to reviews
- Revision(s) submitted
- Revision dates
- Which revisions were reviewed and by which reviewers (anonymized)
- Number of reviewer reports submitted in first round
- Number of revision rounds
- Reviewer interacts with:
  - Editor only - masked
  - Editor only - open
  - Editor and other Reviewer(s) - masked
  - Editor and other Reviewer(s) - open
  - Editor and Author - masked
  - Editor and Author - open
  - Editor, other Reviewer(s) and Author - masked
  - Editor, other Reviewer(s) and Author - open
- Review information published:
  - None
  - Review summaries
  - Review reports
  - Review reports - author opt-in
  - Review reports - reviewer opt-in
  - Submitted Manuscript
  - Submitted Manuscript, author opt-in
  - Editor/Author communication
  - Reviewer identities
  - Reviewer identities, reviewer opt-in
  - Editor identities

#### Quality check(s)

- stats analysis
- RRID checks
- genome checks
- image checks
- plagiarism checks
- COI checks
- reference analysis results
- Technical tools used:
  - Plagiarism checks
  - assess validity/consistency of statistics
  - assess reproducibility/methodological rigor
  - detect image manipulation
  - check references

#### Decision

- Number of editors involved in decision making
- Decision dates
  - Date of acceptance
- Revision
  - Major-revision
  - Minor-revision
- Reject
  - Reject-with-resubmit
- Accept
  - Formal-accept
  - accept-in-principle

#### Publication

- Date of publication
- Timelines/timeliness of editorial process including sent for review
- Version of record: published in journal, proceedings, etc
- Ethics statements
- Funding sources
- Institutions involved

#### Post-publication

- *Events* in Post Publication Peer Review e.g.:
  - requestForReview
  - reviewerAcceptance
  - reviewProduced
- Post publication commenting:
  - Registered commenters
  - Unregistered commenters
  - Editor-selected comments
- *Events* linking to other documents e.g.:
  - Annotations (annotatedBy)
  - linkedTo
- Tweets
- News coverage

## Appendix 4

### Use Cases for DocMaps Framework suggested in Survey 2

READER of a published article: wants to KNOW THE THOROUGHNESS of the peer review process of the article to BETTER DISTINGUISH trustable papers and MAKE A DECISION of further reading/paying to read/citing/sharing/etc.

EDITOR/REVIEWER: wants to MINE reviewer and author comments FROM RELATED published research.

JOURNAL: wants to PUBLISH its own reviews and the reviews transferred from another journal or peer review service alongside the published paper.

PLATFORM: wants to RENDER the publishing process in various visual formats. (e.g. : single merge pdf file with sequence of documents, dates, roles types; separate HTML pages; graphical timeline)

PREPRINT SERVER: wants to IDENTIFY AND SELECT third-party reviews to DISPLAY them alongside the associated preprint.

PREPRINT-FOCUSED EVALUATION AGGREGATION SERVICE that DISPLAYS feeds of events about preprints to BUILD TRUST in that preprint: wants to have an OBJECTIVE LIST of facts about the evolution of a paper (including deposition, quality check, publication and review information) to SUBJECTIVELY INCLUDE these facts in a feed to provide trust indicators to a reader and show the evolution of the paper within the context of its lifetime.

For example: http://hive.review/articles/10.1101/646810?flavour=a shows the context of a review, acceptance, deposition and publication but this was obtained via 3 different services which do not standardise their description of the data. Adding more information here is not possible because a) the cost is too high to build custom integrations which each new event type and b) most other event types are not available in the existing services anyway.

PREPRINT-FOCUSED EVALUATION AGGREGATION SERVICE that DISPLAYS feeds of events about preprints to BUILD TRUST in that preprint: wants a STANDARD way to INGEST peer review content from any provider to easily ADD NEW COMMUNITIES without development effort on either party’s side.

For example: can currently ingest reviews from e.g. Hypothes.is, RSS and custom API integrations, and even standardising the field names across these methods would allow easier onboarding, and faster innovation in the usage of these evaluation events.

PREPRINT-FOCUSED EVALUATION AGGREGATION SERVICE that gives a HOME TO PEER REVIEW COMMUNITIES: wants a STANDARD way to SHOW the editorial and review methods used by a community or on a review so that READERS can easily DETERMINE whether they can apply TRUST to a review based on how it was conducted.

For example: http://hive.review/editorial-communities/316db7d9-88cc-4c26-b386-f067e0f56334 shows Review Commons’ community page and describes their peer review model in free-form text. We could summarise this, filter by it, include it as a badge etc if the terminology were standardised.

AGGREGATOR: wants to FETCH REVIEWS and authors’ replies produced by peer review services and DISPLAY ALONGSIDE the associated preprint and published papers.

AI TEXT MINING APP: wants to crawl and EXTRACT features from referee reports to CLASSIFY, RANK, INDEX papers, preprints, reviewing services, journals.

Generic use case: SOURCE: wants to publicly SHARE as much data related to peer review as possible/desired/allowed in a standard way on the web, so that DESTINATIONS can obtain and process this data and build value-added services.

SOURCE: journal, peer review service, or any other sort of platform that has data relevant to peer review. DESTINATION: interested (authenticated?) parties, including sources.

(More specific use case as a subclass of the above: As a JOURNAL interested in openness, I want to SHARE information about who reviewed what manuscript, when, following what format, with what outcome, and what decision, so that myself and others are ABLE TO ANALYZE, visualize, and otherwise crunch the data.)

## Appendix 5

### Use Case Elements for DocMaps Framework from Meeting 2

#### Use Case: READER/JOURNAL

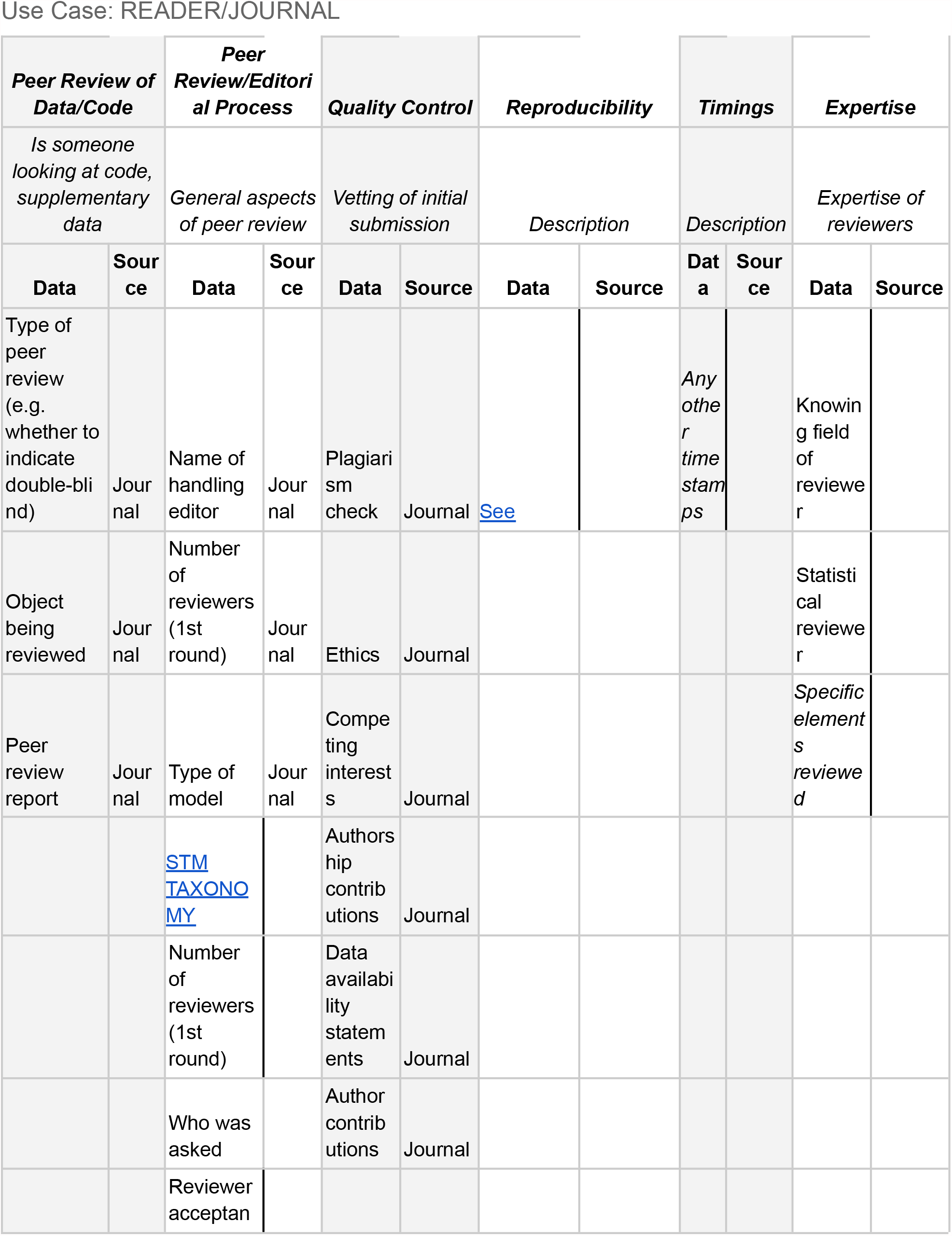
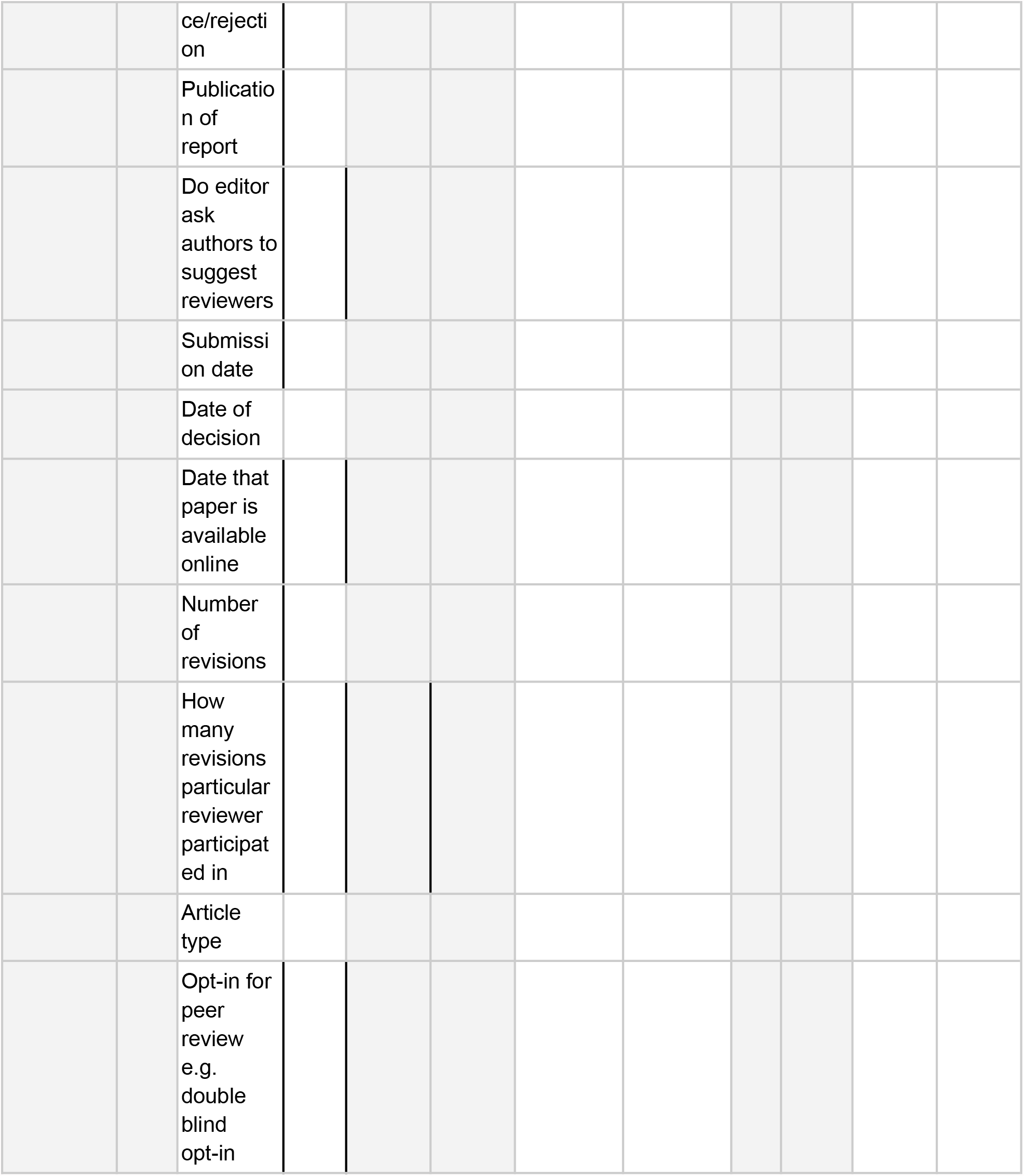

#### Use Case: PREPRINT SERVER

| **Review (Pre-Approved Service)** |  | **Review (Unapproved)** |  | **Non-Review Elements** |  | **Plagiarism Software** |  |
| --- | --- | --- | --- | --- | --- | --- | --- |
| *Description* |  | *Description* |  | *Description* |  | *Description* |  |
| **Data** | **Source** | **Data** | **Source** | **Data** | **Source** | **Data** | **Source** |
| Review ID | (DOI...but may need others) |  |  |  |  |  |  |
| Reviewed Output ID | (DOI...but may need others) |  |  |  |  |  |  |
| License | JATS, others... |  |  |  |  |  |  |
| All of JATS4R Taxonomy | JATS, others... |  |  |  |  |  |  |
| Review Service ID | URI, ROR, others |  |  |  |  |  |  |
| receive_date |  |  |  |  |  |  |  |
| review_accept_date |  |  |  |  |  |  |  |
| review_completion_date |  |  |  |  |  |  |  |
| response_date |  |  |  |  |  |  |  |
| # of reviewers |  |  |  |  |  |  |  |
| reviewer ID | ORCID |  |  |  |  |  |  |
| review report ID |  |  |  |  |  |  |  |
| individual review vs. review report |  |  |  |  |  |  |  |
| structured full text | need more structured version |  |  |  |  |  |  |
| plagiarism score |  |  |  |  |  |  |  |
| Version ID to which review refers |  |  |  |  |  |  |  |
| Score/rating |  |  |  |  |  |  |  |
| reply from authors |  |  |  |  |  |  |  |

## Appendix 6

### The First DocMaps Framework Draft (post-Meeting 2)

The DocMaps Framework

### Summary

We intend DocMaps to be general enough to capture the process behind any document, but our initial focus is capturing evaluative processes surrounding research documents — primarily peer review processes conducted by journals or by services reviewing preprints. We are working with a technical committee of technologists, publishers, preprint server operators, aggregators, and advocates to develop the first use-cases for the framework.

### Core Concepts

#### DocMaps

~~~
{
createdOn: timestamp // The time the DocMap was created
contentType: string // The type of content (book, chapter, review)
content: uri | optional // A link to the content the DocMap refers to
provider: uri | optional // A link identifying the provider of the DocMap
(e.g. the journal)
}
Using just these three pieces of information, a publisher could make a simple assertion that an article exists.
{
createdOn: 07-07-1999T00:00:00Z
contentType: “article”
content: https://doi.org/10.1109/5.771073
provider: https://ieee.org
}
~~~

Of course, this assertion leaves much to be desired. Most DocMaps will contain more information, e.g. the date an article was published; which version of an article the DocMap refers to; *etc*.

DocMaps accomplishes this by building and maintaining schemas for different **Content Types**.

#### Content Types

Although the initial set of content types will be limited, we imagine working with relevant communities to develop many different types that support their use cases. For example: reviews, versions, translations; and discussions.

For now, we’ve defined three different content types with the input of the Technical Committee and the DocMaps core team: **Articles, Reviews**, and **Versions**. For more details, see the draft content type schemas below.

#### Contexts

The real fun starts when you create relationships *between* DocMaps to describe a series of events. These relationships are called **Contexts**, and are defined by a **context key** that describes the type of relationship and the direction of the relationship. Like content types, we imagine that many different types of Contexts will eventually be available, such as:

~~~
// context key: [list of DocMaps]
reviews: [Docmap]
isReviewOf: [Docmap]
versions: [Docmap]
isVersionOf: [Docmap]
translations: [Docmap]
isTranslationOf: [Docmap]
discussions: [Docmap]
isDiscussionOf: [Docmap]
…
~~~

~~~
{
createdOn: 07-07-1999T00:00:00Z
contentType: “article”
content: https://doi.org/10.1109/5.771073
provider: https://ieee.org
contributors: [{name: N. Paskin, type: author}]
title: ‘Toward unique identifiers’
versions: [
          {
          createdOn: 07-07-1999T00:00:Z
          contentType: “version”
          content: https://doi.org/10.1109/5.771073v1
          }
          {
          createdOn: 07-08-1999T00:00:Z
          contentType: “version”
          content: https://doi.org/10.1109/5.771073v2
          }
}
~~~

~~~
{
createdOn: 07-08-1999T00:00:00Z
contentType: “version”
content: https://doi.org/10.1109/5.771073v2
provider: https://ieee.org
isVersionOf: [
       {
       createdOn: 07-07-1999T00:00:Z
       contentType: “article”
       content: https://doi.org/10.1109/5.771073
       }
]
~~~

### Initial Use Cases

#### 1. A publisher captures context about a review of an article published in their journal

~~~
{
contentType: “article”
content: https://doi.org/article/123
createdOn: 2020-01-01T00:00:00Z
provider: https://myjournal.org
title: ‘An article about something!’
contributors: [
          {
          name: “Liz Jones”
          id: https://orcid.org/0002-0002
          role: “author”
          }
          {
          name: “Eric Mays”
          id: https://orcid.org/0005-0001
          role: “data visualization”
          }
]
datePublished: 2020-01-01T:00:00Z
versions: [
          {
          contentType: “version”
          content: https://doi.org/article/123v1
          date_submitted: 2019-12-20T00:00:00Z
          date_online: 2020-08-15T00:00:00Z
          ethics_statements: “This was conducted ethically.”
          competing_interests: “There were no conflicts of interest.”
          }
]
reviews: [
   {
          contentType: “review”
          createdOn: 2020-06-01T00:00:00z
          provider: https://myjournal.org
          decision_date: 2020-07-20T00:00:00z
          decision: ‘accept with revisions’
          contributors:
   [
          {
          name: “John Doe”
          affiliation: “Wassamatta U”
          role: “editor”
            }
          {
          id: 12345
          role: reviewer
          }
          {
             id: 23456
             role: reviewer
          }
          ]
          identity_transparency: ‘double-anonymized’
          reviewer_interacts_with: [editor]
          review_information_published: [editor-identities]
          versions: [
          {
             contentType: “version”
             createdOn: 2020-06-15T00:00:00Z
             contributors: [
          {
          id: 12345
          role: reviewer
          }
          {
          id: 23456
          role: reviewer
          }
          ]
          }
          {
             contentType: “version”
             createdOn: 2020-07-10T00:00:00Z
             contributors: [
          {
          id: 12345
          role: reviewer
          }
         ]
       }
     ]
   }
  }
~~~

### Data Elements

| ***Name*** | ***Value*** | ***DocMaps Location*** |
| --- | --- | --- |
| Submission date | 2019-12-20 | article -> versions -> date_submitted |
| Date of decision | 2020-07-20 | article -> reviews -> decision_date |
| Date that paper is available online | 2020-08-15 | article -> versions -> date_online |
| Publication of report | none | article -> reviews -> (no content field provided) |
| Decision / Score | accept with revisions | article -> reviews -> decision |
| Number of reviewers (1st round) | 2 | article -> reviews -> first version -> count of contributors with role "reviewer" |
| Publication of response | none | article -> reviews -> (no author response provided) |
| Review info published | editor identities | article -> reviews -> review_information_published |
| Who was asked | 12345, 23456 | article -> reviews -> count of contributors with roles "reviewer" or "invited_reviewer" |
| Do editors ask authors to suggest reviewers | no | article -> reviews -> contributors (no author_suggested_reviewer on any contributors) |
| Name of handling editor | John Doe | article -> reviews -> contributors (editor) |
| Type of model | double anonymized | article -> reviews -> identity_transparency |
| Opt-in for peer review<br>e.g. double blind opt-in | no | article -> reviews -><br>identity_transparency<br>(opt-in not provided<br>from STM schema) |
| Number of revisions | 2 | article -> reviews -><br>versions |
| How many revisions<br>particular reviewer<br>participated in | 12345: 2<br>23456: 1 | article -> reviews -><br>versions -><br>contributors |
| Reviewer interacts<br>with | editor | article -> reviews -><br>reviewer_interacts_wit<br>h |
| Post publication<br>commenting | none | article -> reviews -><br>(no<br>post_publication_comm<br>enting provided) |
| Object type | article | article -> contentType |
| Ethics | This was conducted<br>ethically. | article -> version -><br>ethics_statements |
| Competing interests | There were no conflicts<br>of interest | article -> version -><br>competing_interests |
| Data availability<br>statements | none | article -> version -> (no<br>data_availability_state<br>ments) provided |
| Authorship<br>contributions | Liz Jones, author<br>Eric Mays, data<br>visualization | article -> contributors |

#### 2. An independent review service notifies a preprint server about a review of an article on their platform

~~~
{
contentType: “review”
content: https://doi.org/review/123
createdOn: 2020-01-01T00:00:00z
provider: https://myreviewservice.org
decision_date: 2020-07-20T00:00:00z
decision: “accept”
contributors: [
          {
          name: “Tricia McMillan”
          affiliation: “Maximegalon University”
          role: “editor”
          id: https://orcid.org/0000-0000
          author_suggested: false
          },
          {
          name: “Zaphod Beeblebrox”
          affiliation: “Betelgeuse State College”
          role: reviewer
          id: https://orcid.org/0001-0001
          author_suggested: true
          }
          {
          name: “Arthur Dent”
          affiliation: “BBC” role: reviewer
          id: https://orcid.org/0002-0002
          }
          {
          name: “Ford Prefect”
          affiliation: “Pan Galactic Gargle Blaster Society”
          role: invited_reviewer
          id: https://orcid.org/0002-0002
          }
]
author_response: https://doi.org/response/123
identity_transparency: [all-identities-visible, opt-in]
reviewer_interacts_with: [editors, reviewers, authors]
review_information_published: [reviewer-identities, editor-identities,
review-reports-author-opt-in]
versions: [
          {
          contentType: “version”
          date_submitted: 2020-06-15T00:00:00Z
          }
]
isReviewOf: [
          {
          contentType: “article”
          content: https://doi.org/preprint/123
          }
]
~~~

| ***Name*** | ***Value*** | ***DocMaps Location*** |
| --- | --- | --- |
| Submission date | 2020-06-15 | review -> versions -> date_submitted |
| Date of decision | 2020-07-20 | review -> decision_date |
| Date that paper is available online |  | n/a for this example, data can be obtained from preprint server or crossref |
| Publication of report | Yes<br>(https://doi.org/review/123) | review -> content |
| Decision / Score | accept | review -> decision |
| Number of reviewers (1st round) | 2 | review -> count of contributors with role "reviewer" |
| Publication of response | Yes<br>(https://doi.org/response/123) | review -> author_response |
| Review info published | reviewer-identities, editor-identities, review-reports | review -> review_information_published |
| Who was asked | Zaphod Beeblebrox, Arthur Dent, Ford Prefect | article -> reviews -> count contributors with roles "reviewer" or "invited_reviewer" |
| Do editor ask authors to suggest reviewers | yes | article -> reviews -> contributors -> author_suggested: true (Zaphod Beeblebrox) |
| Name of handling editor | Tricia McMillan | review -> contributors (editor) |
| Type of model | fully transparent | article -> reviews -> identity_transparency |
| **Name** | **Value** | **DocMaps Location** |
| Opt-in for peer review<br>e.g. double blind opt-in | yes | article -> reviews -><br>identity_transparency |
| Number of revisions | 2 | article -> reviews -><br>versions |
| How many revisions<br>particular reviewer<br>participated in | Arthur Dent: 1<br>Tricia McMillan: 1<br>Zaphod Beeblebrox: 1<br>Ford Prefect: 0 | implied by convention<br>from review -> version<br>with no reviewers<br>specified, but<br>reviewers specified at<br>review -> contributors<br>level |
| Reviewer interacts<br>with | editors, reviewers,<br>authors | review -><br>author_interacts_with |
| Post publication<br>commenting | none | article -> reviews -><br>(no<br>post_publication_comm<br>enting provided) |
| Object type | review | review -> contentType |
| Ethics |  | n/a for this example,<br>data can be obtained<br>from preprint server or<br>crossref |
| Competing interests |  | n/a for this example,<br>data can be obtained<br>from preprint server or<br>crossref |
| Data availability<br>statements |  | n/a for this example,<br>data can be obtained<br>from preprint server or<br>crossref |
| Authorship<br>contributions | Tricia McMillan, editor<br>Arthur Dent, reviewer | review -> contributors |
|  | Zaphod Beeblebrox,<br>Reviewer |  |

### Draft Content Type Schemas

#### Article

~~~
{
// Common DocMap fields
contentType: “article”
content: uri // A canonical link or DOI to the article, if it exists
createdOn: timestamp
provider: uri
// Article specific metadata
title: string
abstract: string contributors:
[Contributors] datePublished: date
// Contexts
versions: [Version]
}
~~~

#### Review

~~~
{
// Common DocMap fields
contentType: “review”
content: uri // A canonical link or DOI to the report publication, if it
exists
createdOn: timestamp
provider: uri
// Review specific metadata fields
decision_date: timestamp
decision: string
contributors: [Contributor] // Includes reviewers, editors, asked reviewers
author_response: uri
identity_transparency: string // STM Assoc taxonomy
reviewer_interacts_with: [string] // STM Assoc taxonomy
review_information_published: [string] // STM Assoc taxonomy
post_publication_commenting: [string] // STM Assoc taxonomy
// Contexts
isReviewOf: [Article] // A list of DocMaps of reviewed objects
}
~~~

#### Version

~~~
{
// Common DocMap fields
contentType: “version”
content: uri // A canonical link or DOI to the version of the referenced
object, if it exists
createdOn: timestamp
provider: uri
// Version specific metadata
date_submitted: timestamp date_online: timestamp
ethics_statements: string
competing_interests: string
data_availability: string
contributors: [Contributor]
}
~~~

## Appendix 7

### Alignment of JATS4R and DocMaps elements

#### Use Case 1: A publisher captures context about a review of an article published in their journal

| JATS4R Element | Value | DocMaps Element |
| --- | --- | --- |
| sub-article | n/a | article -> reviews |
| front-stub | n/a | article -> reviews -> {...} |
| article-id | https://doi.org/review/abcd | article -> reviews -> [content] |
| contrib /<br>contrib-group | John Doe, 12345, 23456 | article -> reviews -> [contributors] |
| contrib-type | John Doe (author) | mapped from article -> reviews -> [contributors -> role] |
| anonymous | 12345, 23456 | implied via article -> reviews -> [contributors (no name or orcid)] |
| role -><br>@specific-use | John Doe (editor), 12345 (referee <- referee), 23456 (referee <- referee) | mapped from mapped from article -> reviews -> [contributors -> role] |
| permissions | n/a | n/a (but we should add this) |
| pub-date | 2020-08-15T00:00:00Z | article -> reviews -> version [1] -> date_online |
| related-object | 2 versions | article -> reviews -> versions (or any nested DocMap) |
| related-object -> @document-type | potentially not enough info to map from DocMaps content types | article -> reviews -> versions (or any nested DocMap) -> contentType<br>OR<br>article -> reviews -> contentType when nested DocMaps exist |
| related-object -> @source-id | https://myjournal.org | article -> reviews -> provider |
| pub-history / event | 2 referee-report-received, 1 editor-decision-sent | article -> reviews -> versions (or any nested DocMap) |
| pub history / event -> @event-type | round 1 referee-report-received: 2020-06-15T00:00:00Z<br>round 2 referee-report-received: 2020-07-10T00:00:00Z<br>editor_decision_sent: 2020-07-20T00:00:00z | round 1 mapped from article -> reviews -> versions[0] -> createdOn<br>round 2 mapped from article -> reviews -> versions[1] -> createdOn<br>editor decision mapped from article -> reviews -> decisionDate |
| institution | none | article -> reviews -> contributors -> affiliation |
| competing interests | none | article -> reviews -> competing_interests |
| referee_number | 12345: 1<br>23456: 2 | implied from order of array: article -> reviews -> contributors |
| peer_review_stage | pre-publication | implied from review date_online < article date_online |
| transfer | no | implied from provider being same value on both article and article -> reviews |
| revision_round | 2 | implied from number of elements in article -> reviews -> versions |
| recommendation | accept <- accept with revisions | mapped from article -> reviews[0] -> decision |
| peer_review_type | double-blind <- double-anonymized | mapped from article -> reviews[0] -> identity_transparency (uses STM taxonomy) |

#### Use Case 2: An independent review service notifies a preprint server about a review of an article on their platform

Use Case 2: An independent review service notifies a preprint server about a review of an article on their platform.
| JATS4R Element | Value | DocMaps Element |
| --- | --- | --- |
| article | n/a | review |
| article-metadata | n/a | review -> {...} |
| article-id | https://doi.org/review/123 | review -> content |
| contrib /<br>contrib-group |  | reviews -> [contributors] |
| contrib-type | Tricia McMillan (author) | mapped from article -> reviews -> [contributors<br>-> role] |
| anonymous | 12345, 23456 | implied via article -> reviews -> [contributors (no<br>name or orcid)] |
| role -><br>@specific-use | Tricia McMillan (editor),<br>Zaphod Beeblebrox (referee<br><- reviewer), Athur Dent<br>(referee <- reviewer) | mapped from mapped from article -> reviews -><br>[contributors -> role] |
| permissions | n/a | n/a (but we should add this) |
| pub-date | 2020-07-31T00:00:00Z | review -> versions [0] -> date_online |
| related-object | 1 article | review -> isReviewOf [0] |
| related-object -> @document-type | peer-reviewed-article <- article | review -> isReviewOf [0] -> contentType |
| related-object -> @source-id | https://myreviewservice.org | article -> reviews -> provider |
| pub-history / event | 1 referee-report-received, 1 editor-decision-sent, 1 author-comment-received | article -> reviews -> versions (or any nested DocMap) |
| pub history / event -> @event-type | 1 referee-report-received: 2020-07-15T00:00:00Z<br>editor_decision_sent: 2020-07-20T00:00:00z<br>author-comment-received: 2020-07-15T00:00:00Z | review -> versions[0] -> date_submitted<br>review -> decisionDate<br>review -> author_comment -> date_submitted |
| institution | Tricia McMillan (University of Maximegallon)<br>Zaphod Beeblebrox (Beetelgeuse State College)<br>Arthur Dent (BBC) | article -> reviews -> contributors -> affiliation |
| competing interests | none | article -> reviews -> competing_interests |
| referee_number | Zaphod Beebelbrox: 1<br>Arthur Dent: 2 | implied from order of array: article -> reviews -> contributors |
| peer_review_stage | n/a | unknown, as article being reviewed is only referenced by ID. Could maybe be implied as "post-publication" from lack of provider or date_online for isReviewOf docmap |
| transfer | n/a | unknown, as article being reviewed is only referenced by ID. could maybe be implied as "yes" from lack of provider for isReviewOf docmap |
| revision_round | 1 | implied from number of elements in review -> versions |
| recommendation | accept -> accept | mapped from review -> decision |
| peer_review_type | visible -><br>all_identities_visible, opt-in | mapped from review -> identity_transparency (uses STM taxonomy) |

